# Capping protein insulates Arp2/3-assembled actin patches from formins

**DOI:** 10.1101/647198

**Authors:** Ingrid Billault-Chaumartin, Sophie G. Martin

## Abstract

How actin structures of distinct identities and functions co-exist within the same environment is a critical self-organization question. Fission yeast cells have a simple actin cytoskeleton made of four structures: Arp2/3 assembles actin patches around endocytic pits; the formins For3, Cdc12 and Fus1 assemble actin cables, the cytokinetic ring during division, and the fusion focus during sexual reproduction, respectively. The focus concentrates the delivery of hydrolases by myosin V to digest the cell wall for cell fusion. We discovered that cells lacking capping protein (CP), a heterodimer that blocks barbed-end dynamics and associates with actin patches, exhibit a delay in fusion. Consistent with CP-formin competition for barbed-end binding, Fus1, F-actin and the linear filament marker tropomyosin hyper-accumulate at the fusion focus in absence of CP. However, myosin V and exocytic cargoes are diverted to ectopic foci and reduced at the fusion focus, which underlies the fusion defect. Remarkably, ectopic foci coincide with actin patches, which now contain low levels of Fus1. During mitotic growth, actin patches lacking CP similarly display a dual identity, as they accumulate the formins For3 and Cdc12 and are co-decorated by tropomyosin and the patch marker fimbrin. Thus, CP serves to protect Arp2/3-nucleated structures from formin activity.

## Introduction

Cells simultaneously contain several actin-based structures that need to be tailored to their specific function, with a specific architecture, size, life-time, and set of actin binding proteins. The specific architecture is defined in part by the nucleator that assembles it [1, 2]: Arp2/3 promotes the assembly of branched structures, whereas other nucleators, in particular formins, assemble linear ones. The dendritic networks assembled by Arp2/3 generate pushing forces against membranes, for instance in the lamellipodium of migrating cells, to drive the movement of intracellular bacteria, or to promote internalization of endocytic vesicles in yeast actin patches. Formin-nucleated actin structures consist of linear filaments, which can be bundled in parallel or antiparallel manner for protrusive or contractile force generation, for instance in filopodia or cytokinetic contractile ring, respectively, or which underlie long-range myosin-based transport.

The principles that allow a cell to assemble distinct functional actin structures at the same time are just beginning to be understood. First, because the building blocks for assembly of diverse actin structures are the same, competition is an important factor. For instance, diverse filamentous actin (F-actin) structures are in competition for a limited pool of actin monomers [3]. This competition is modulated by profilin associated with G-actin, which favors F-actin assembly by formins and other nucleators over Arp2/3 [4, 5]. Second, the structure’s identity may be conferred, at least in part, by the specific actin nucleator that assembles it. For instance, formins promote the formation of more flexible filaments [6], which favor tropomyosin association [7]. Different formins were recently proposed to promote association of distinct tropomyosin isoforms on filaments [8]. Third, self-assembly principles likely govern the segregation of specific actin-binding proteins to diverse structures. For instance, competition between fimbrin and tropomyosin for F-actin binding, together with their individual cooperative loading on actin filament, drives their association to distinct actin structures [9]. This is manifested in vivo by preferential association of tropomyosin to formin-assembled structures and fimbrin to Arp2/3-nucleated actin patches [10, 11].

An important factor that helps limit actin filament growth is capping protein (CP). CP is present in cells in µM concentration, similar to the concentration of actin filament barbed ends, and binds the barbed end to arrest dynamics [12]. CP forms a heterodimer of structurally similar α- and β-subunits, both of which harbour a mobile C-terminal extension, called the tentacle, which strongly contributes to barbed end binding [13–15]. CP lacking both tentacles still forms a stable complex but does not bind actin, with the α-tentacle playing a more critical role than the β-tentacle, both in vitro and vivo [14, 16]. CP activity is further regulated by interaction with binding partners bearing a capping protein interaction (CPI) motif [17]. Indeed, CP carrying surface mutations that block CPI motif binding retain capping activity in vitro, but lose localization and function in vivo, indicating that binding partners are required for its activity in vivo [18]. Furthermore, the Aim21/Tda2 complex, which binds CP through the same surface residues, modulates CP recruitment and activity at actin patches in yeast [19, 20]. By keeping filaments short, CP promotes Arp2/3 branching and plays a major role in the force production of dendritic networks [21]. Indeed, absence of CP in vivo leads to loss of the lamellipodium in migrating cells [22, 23], and excess actin filaments in yeast actin patches, which exhibit a longer lifetime [16, 24–26].

As formins and capping proteins both interact with the barbed end of actin filaments but promote opposite activities – i.e. filament extension vs. capping – they compete with each other in vitro [27–30]. Interestingly, recent single-molecule work has shown that formin and CP can simultaneously bind the filament barbed end, forming a ternary ‘decision complex’ intermediate [31, 32]. Evidence for competition in vivo comes from work in the fission yeast *Schizosaccharomyces pombe*, in which deletion of capping protein ameliorates the function of a hypomorphic formin *cdc12* allele for cell division [33]. CP may also compete with formins and ENA/VASP to control filopodial shape [34]. Whether such competition contributes to actin structure identity has not been explored.

The fission yeast cell represents a simple system in which to dissect the mechanisms by which distinct actin structures are formed. Indeed, this cell contains only four actin structures, actin patches, cables, ring and focus, each of which fulfils a specific function [1, 35]. Arp2/3 nucleates the assembly of actin patches around invaginating endocytic vesicles. The actin patch, marked in particular by fimbrin Fim1, reproducibly assembles in a stereotypical manner [36, 37] and is thought to provide force for vesicle internalization [38]. CP localizes to actin patches [33, 39, 40], where it limits actin incorporation and helps force production [16, 24–26]. Three formins nucleate distinct structures formed of linear filaments: Cdc12 promotes the assembly of the actin contractile ring for cell division; For3 assembles linear actin filaments bundled in cables that underlie long-range myosin-based transport for polarized cell growth; Fus1 is expressed specifically during cell mating and nucleates the assembly of the fusion focus, an aster-like actin structures that concentrates secretory vesicles transported by the myosin V Myo52 [41, 42]. These vesicles carry cell wall hydrolases such as Agn2 and Eng2, whose local secretion drives cell wall digestion for cell fusion. Besides Fus1, the coalescence of the fusion focus requires the action of tropomyosin Cdc8 [43], and a visual screen revealed Cdc8 functions together with the type V myosin Myo51 and associated proteins to organize the focus in a single structure [44]. This same screen identified the deletion of *acp2* to have fusion focus defects.

Starting from the hypothesis that CP and formin Fus1 compete during fusion focus assembly, we discovered that CP protects actin patches against formin activity. In the absence of CP, formin Fus1 binds uncapped barbed ends in actin patches, forming ectopic foci that divert secretory vesicles away from the site of cell-cell contact and compromise cell fusion. Similarly, in proliferating cells lacking CP, the formins For3 and Cdc12 are ectopically recruited to actin patches, which exhibit a dual identity manifested by co-decoration with fimbrin and tropomyosin. Thus, CP ensures actin structure identity by insulating Arp2/3-assembled structures against formins.

## Results

### Capping protein is required for efficient cell-cell fusion

To investigate the role of capping proteins (CP) in cell fusion, we first assessed the fusion efficiency of strains lacking one or both CP subunits (*acp1*Δ, *acp2*Δ, *acp1*Δ *acp2*Δ). After 12 hours of starvation, these strains exhibited a reduced fraction of fused zygotes compared to WT, which increased 36 hours post starvation (Figure 1A), indicating that the absence of CP causes a cell fusion delay. We measured the duration of the fusion process, from initial formation of the fusion focus marked by the type V myosin Myo52 in both partner cells [41] to cytoplasmic mixing. Cytoplasmic mixing was defined by entry in the M-cell of cytosolic GFP expressed in P-cells under control of the *p*^*map3*^ promoter. The process lasted significantly longer in CP lacking cells (Figure 1B-C). We also observed that the fusion focus persisted significantly longer post-fusion in CP lacking cells (Figure 1D-E). Both phenotypes were clear in all mutant combinations, but strongest in *acp2*Δ single mutant, on which we focused most of our attention. Thus, CP promotes the fusion process, since its absence causes fusion delay and persistence of the fusion focus after fusion.

**Figure 1.**
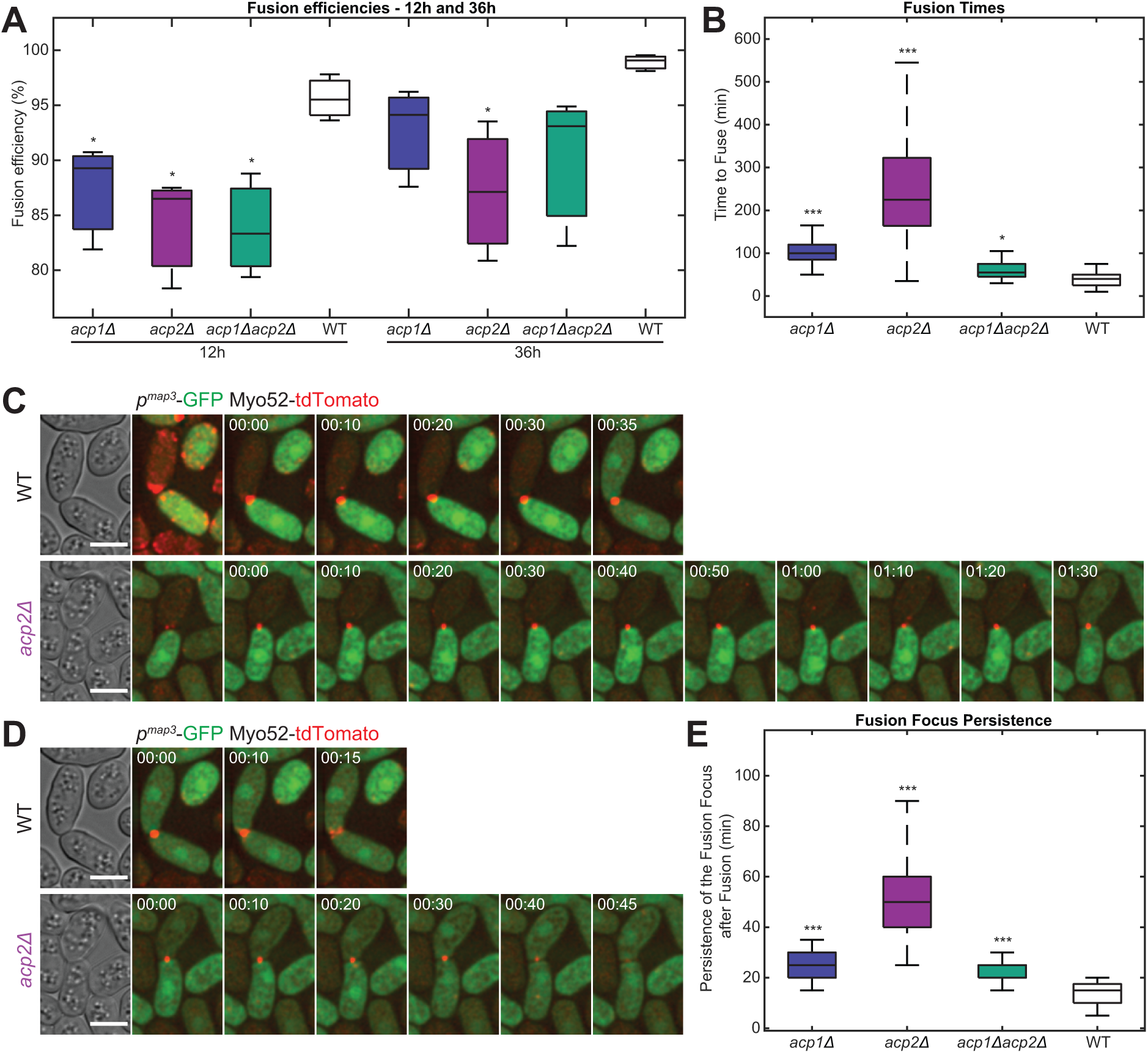
Absence of CP leads to fusion delay and persistence of the fusion focus after fusion. **A.** Boxplot of fusion efficiencies 12h and 36h after nitrogen removal in *acp1Δ, acp2Δ, acp1Δ acp2Δ* and WT strains. p-values relative to WT are 4.9×10^−2^, 2.1×10^−2^ and 1.7×10^−2^ for *acp1Δ, acp2Δ* and *acp1Δacp2Δ* at 12h, and 7.6×10^−2^, 3.3×10^−2^ and 9.1×10^−2^ at 36h, respectively. Boxplots are made on 3 replicates where ~250 mating pairs per condition were quantified for each replicate. **B.** Boxplot of fusion times in *acp1Δ, acp2Δ, acp1Δ acp2Δ* and WT strains. p-values relative to WT are 3.3×10^−17^, 2.2×10^−11^ and 8.3×10^−4^ for *acp1Δ, acp2Δ* and *acp1Δacp2Δ*, respectively. This was measured on 32, 53, 97 and 56 individual mating pairs for WT, *acp1Δ, acp2Δ* and *acp1Δacp2Δ*, respectively. **C.** Time-lapse images of Myo52-tdTomato and cytosolic GFP expressed in P-cells under *map3* promoter in WT and *acp2Δ*, from beginning to end of the fusion process. The beginning is defined as the first instance of the focus in both cells, the end as entry of the GFP into the M-cell cytosol. **D.** Time-lapse images of strains as in (C) from fusion time to disappearance of the fusion focus. **E.** Boxplot of fusion focus persistence times in *acp1Δ, acp2Δ, acp1Δacp2Δ* and WT. p-values relative to WT are 2.7×10^−15^, 2.4×10^−22^ and 8.1×10^−10^ for *acp1Δ, acp2Δ* and *acp1Δacp2Δ*, respectively. This was measured on 32, 53, 97 and 56 individual mating pairs for WT, *acp1Δ, acp2Δ* and *acp1Δacp2Δ*, respectively. Time in hour:min. Bars are 5µm.

### Formin Fus1 and F-actin excessively accumulate at the fusion site in absence of capping protein

Consistent with CP preventing filament barbed-end extension, previous work reported that Arp2/3-assembled actin patches lacking CP accumulate more actin [24, 40]. To investigate the organization of F-actin during fusion, we first used GFP-CHD as general F-actin marker [45, 46]. Like in interphase cells, actin patches appeared brighter in *acp2*Δ than WT cells during mating (Figure 2A) [24]. We also observed significantly more F-actin at the position of the fusion focus in *acp2*Δ, *acp1Δ* and *acp1Δ acp2Δ* than WT cells (Figure 2A, D; Figure S1A). Because GFP-CHD fluorescence intensity measurements at the site of cell-cell contact cannot distinguish between F-actin incorporation in the fusion focus or in surrounding patches, we probed the localization of specific fusion focus components. The formin Fus1 accumulated approximately 4-fold more in *acp2Δ, acp1Δ* and *acp1Δ acp2Δ* cells than in WT cells at the fusion focus (Figure 2B, D; Figure S1B). We also noted a mild increase in global Fus1-sfGFP fluorescence levels (Figure S1D). Likewise, tropomyosin Cdc8, which preferentially binds formin-assembled linear actin filaments [7, 43, 47–49], showed about 2-fold increased levels at the fusion focus of *acp2Δ* compared to WT cells (Figure 2C-D; Figure S1C). Thus, similar to previous observations for actin patches, the absence of capping protein leads to increased F-actin and associated proteins at the fusion focus.

**Figure 2.**
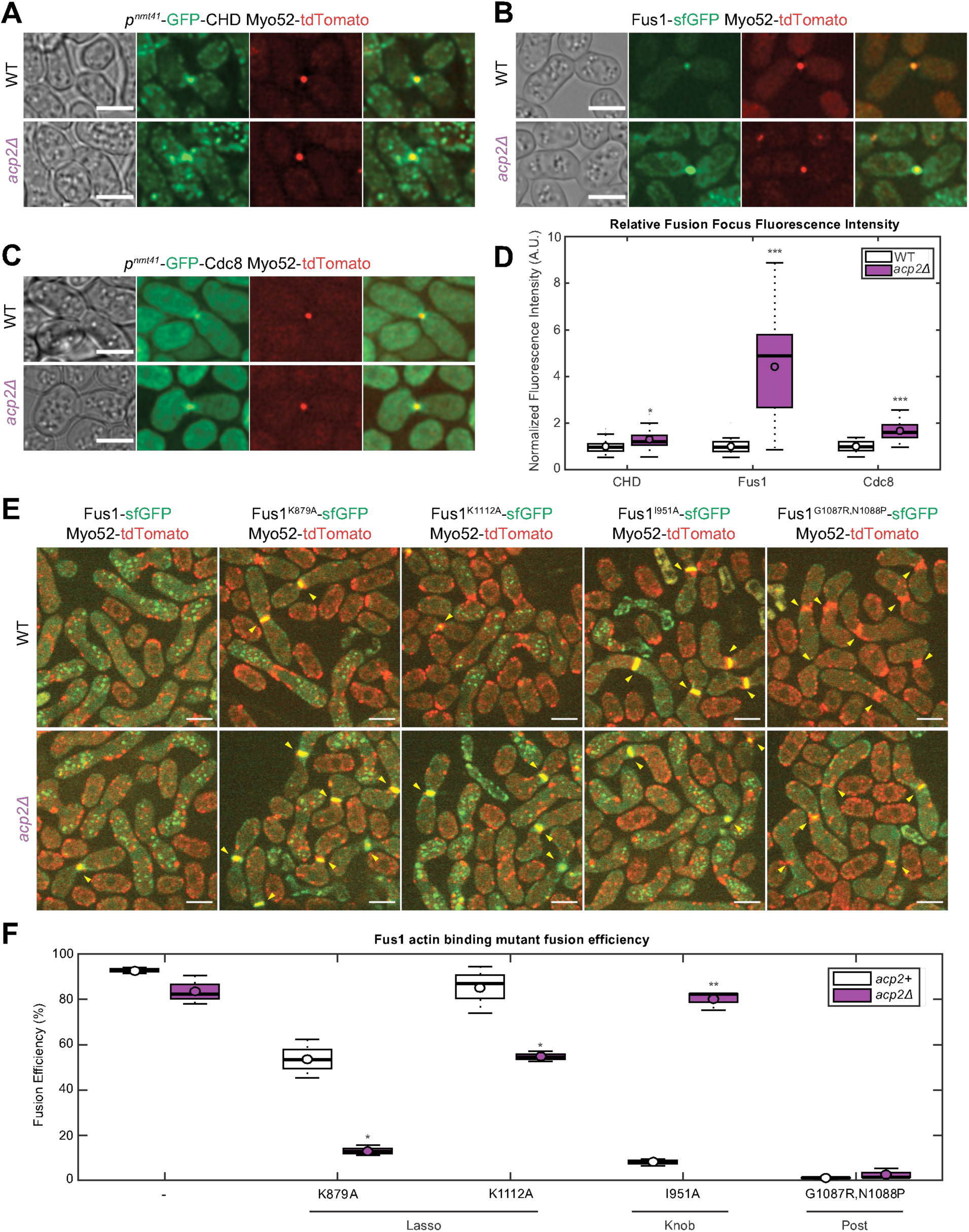
Absence of CP leads to increased actin, tropomyosin and Fus1 at the fusion focus. **A-C.** Myo52-tdTomato and (A) GFP-CHD labeling F-actin, (B) Fus1-sfGFP and (C) GFP-Cdc8 in WT and *acp2Δ* at fusion time. **D.** Boxplot of fusion focus fluorescence intensities normalized to WT at fusion time in strains as in (A-C). p-values relative to WT are 2.9×10^−4^, 2.0×10^−9^ and 1.3×10^−8^ for CHD, Fus1 and Cdc8, respectively (for both strains n = 40, 24 and 24 individual mating pairs, respectively). See Figure S1A-C for full fluorescence profile measurements. **E.** Spinning-disk confocal microscopy images of Myo52-tdTomato and WT or mutant Fus1-sfGFP, as indicated, in WT and *acp2Δ*. Arrowheads point to unfused cell pairs, which exhibit a broad distribution of Fus1 and Myo52 at the zone of contact. **F.** Boxplot of fusion efficiencies at 9h after nitrogen removal in WT or *acp2Δ* strains carrying WT or mutant *fus1*, as indicated. p-values of the *acp2Δ* strain relative to the corresponding WT are 7.2×10^−2^, 1.3×10^−3^, 8.1×10^−3^, 1.0×10^−5^ and 2.8×10^−1^ for *fus1, fus1*^*K879A*^, *fus1*^*K1112A*^, *fus1*^*I951A*^ *and fus1*^*GN1087,1088RP*^, respectively. Boxplots are made on 3 replicates where ~450 mating pairs per strain were quantified for each replicate. Bars are 5µm.

**Figure S1 – related to Figure 2.**
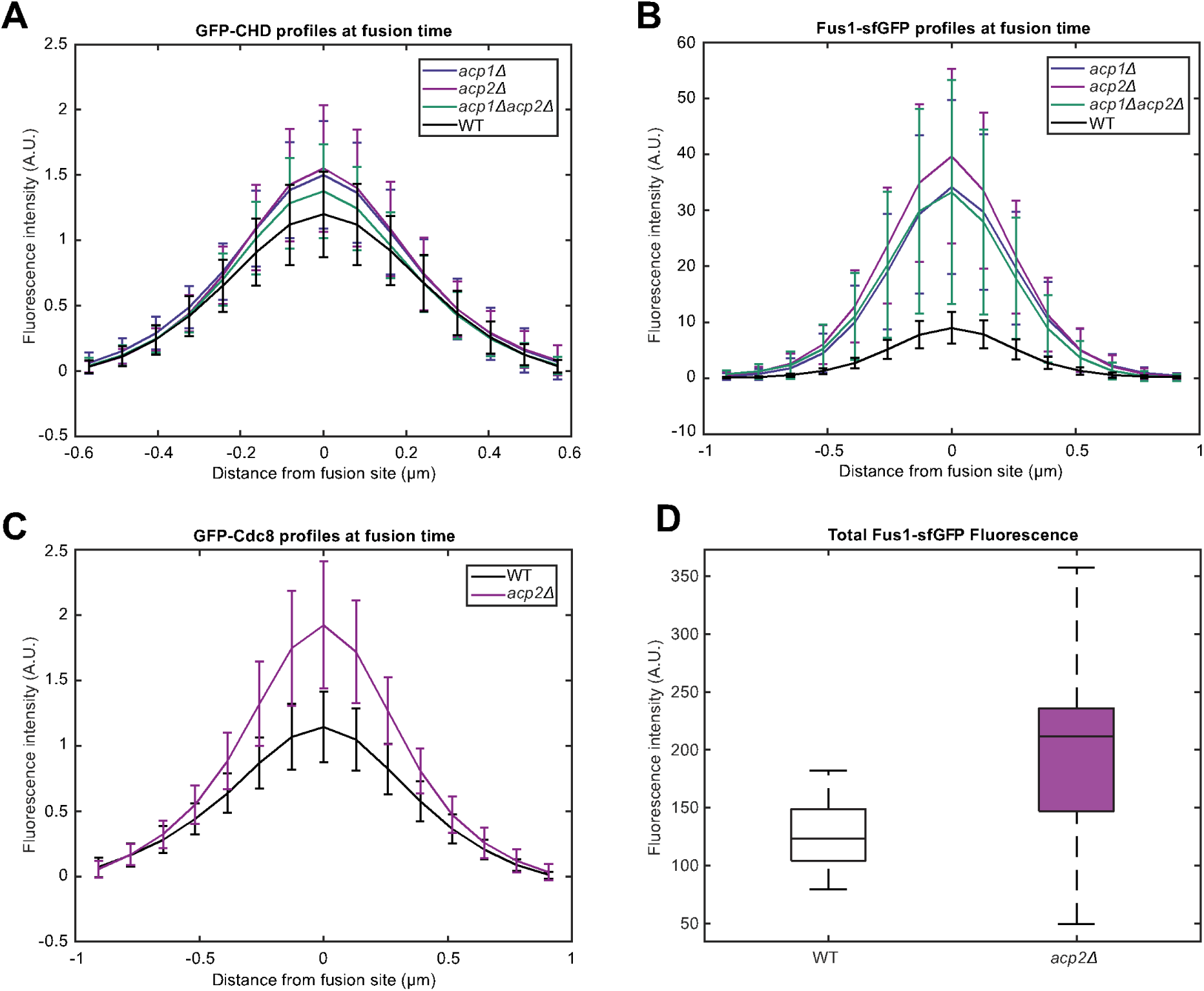
Quantification of the increase in actin, tropomyosin and Fus1 at the fusion focus in absence of CP. **A-C.** Profiles of the bleach-corrected fluorescence intensities around the fusion focus at fusion time in the strains shown in Figure 2. The boxplot in Figure 2D shows the central points of these profiles further normalized to WT. (A) GFP-CHD profiles. (B) Fus1-sfGFP profiles (C) GFP-Cdc8 profiles. **D.** Boxplot of total Fus1 fluorescence intensity in fusing cells, in WT and *acp2Δ*. p-value relative to WT is 6.7×10^−2^. This was measured on 33 and 55 individual mating pairs for WT and *acp2Δ* cells, respectively.

**Figure S2 – related to Figure 2.**
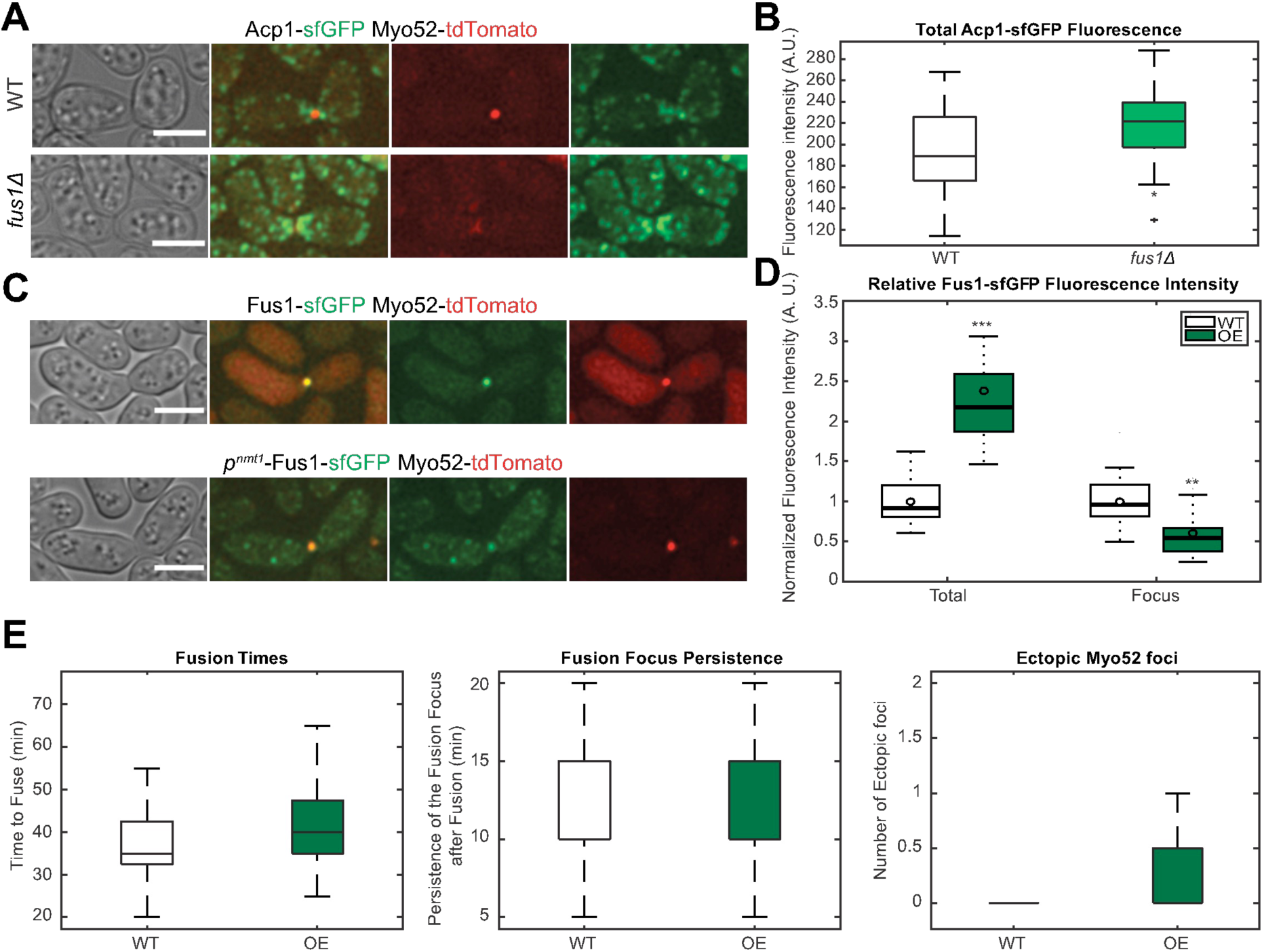
Weak increase in CP fluorescence in *fus1*Δ and no effect of Fus1 overexpression. **A.** Myo52-tdTomato and Acp1-sfGFP in WT and *fus1*Δ at fusion time. **B.** Boxplot of total fluorescence intensities in fusing cells in strains as in (A). p-value relative to WT is 1.4×10^−4^ (n = 51 individual mating pairs for each strain). **C.** Myo52-tdTomato and Fus1-sfGFP at fusion time, in strains expressing Fus1 either from endogenous locus or, in addition, under the *nmt1* promoter. **D.** Boxplot of total and fusion focus fluorescence intensities normalized to WT in fusing cells in strains as in (C). p-value relative to WT is 1.3×10^−12^ for the total intensity (n = 15 and 45 mating pairs for WT and over-expressing cells) and 3.2×10^−6^ for the fusion focus intensity (n = 30 mating pairs each). Note that the quantified levels of overexpressed Fus1-sfGFP do not represent all Fus1 in the cell, as endogenous Fus1 is not tagged. **E.** Boxplots of fusion times, fusion focus persistence, and ectopic Myo52 foci in WT and Fus1-overexpressing strains. p-values relative to WT are 5.2×10^−2^ for fusion times (n = 60), 4.2×10^−1^ for persistence times (n = 60) and 3.7×10^−1^ for ectopic foci (n = 44 and 36 individual cells for WT and Fus1-overexpressing cells, respectively). Bars are 5µm.

The dramatic increase in Fus1 levels in *acp2*Δ is consistent with the proposed competition between formins and CP for barbed end binding [31–33]. We note that, conversely, CP levels were mildly increased in *fus1Δ* (Figure S2A-B). To probe the hypothesis that CP promotes cell fusion by limiting Fus1-driven actin polymerization at the fusion focus, we tested whether 1) Fus1 overexpression mimics loss of CP, 2) reducing Fus1 activity ameliorates fusion efficiency in absence of CP, and 3) CP localizes to the fusion focus. Overexpressing Fus1 did not change the duration of the fusion process, the persistence of the fusion focus, or the presence of ectopic Myo52 foci (Figure S2C-E; see below regarding ectopic foci). However, while overexpression increased the total Fus1 levels, it did not increase Fus1 levels at the fusion focus as observed in *acp2*Δ (Figure S2D). To reduce Fus1 activity, we constructed four Fus1 alleles mutated in the FH2 domain. While all four mutants completely abolished actin assembly in vitro [50], they showed differential cellular phenotypes when introduced as sole copy at the native genomic locus of otherwise wildtype cells. Fus1^K879A^ and Fus1^K1112A^, which carry mutations in the FH2 lasso, were partly fusion-competent, whereas Fus1^I951A^ and Fus1^G1087R,N1088P^, which carry mutations in the FH2 knob and post, respectively, almost completely blocked cell fusion. Combining these *fus1* alleles with *acp2Δ* phenotype did not systematically ameliorate the fusion phenotype (Figure 2E-F). In particular, the two hypomorphic lasso mutants compromised fusion further in absence of *acp2*. By contrast, the almost completely fusion-incompetent *fus1*^*I951A*^ allele permitted high levels of cell fusion upon *acp2* deletion. This reveals an allele-specific suppression, where only Fus1 knob, but not lasso mutants compromise the competition with CP. This finding is consistent with the recently proposed steric clash between the FH2 knob and the CP*β* tentacle [31]. However, this allele-specific competition may principally take place elsewhere than the fusion focus because neither Acp1 nor Acp2 were detected at this location (see Figure 5A, C, D, F).

### Absence of capping protein leads to reduced levels of myosin V and cargoes at the fusion focus and formation of ectopic foci

Unexpectedly, in contrast to actin and Fus1, the type V myosin Myo52 was reduced about 2-fold at the fusion focus in *acp2Δ* cells compared to WT cells (Figures 3F; see also Figure 1C). A similar reduction was observed in *acp1*Δ and *acp1*Δ *acp2*Δ cells (Figure S3A). Since Myo52 brings the glucanase-containing exocytic vesicles to the region of cell contact, we further probed the accumulation of exocytic markers. The exocyst subunits Exo84 and Exo70, the Rab11-homologue Ypt3, and the glucanases Agn2 and Eng2, all displayed a reduced intensity at the fusion focus at fusion time in CP-lacking strains (Figure 3A-F; Figure S3B-F). Since these glucanases are directly responsible for degrading the cell wall between the two cells in contact [41], it is likely that their reduced accumulation at the fusion focus is the cause of the *acp2Δ* fusion delay.

**Figure 3.**
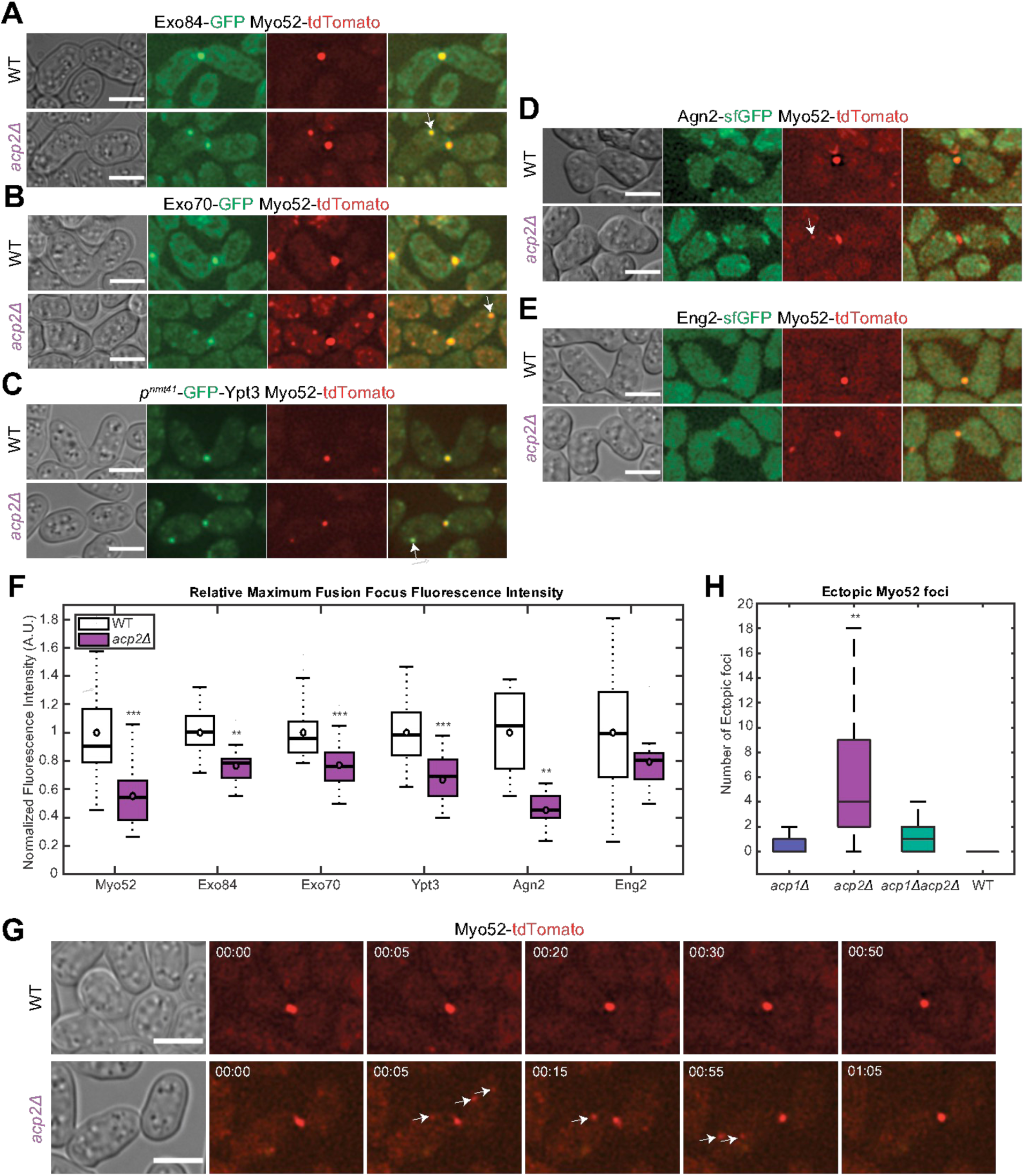
Absence of CP leads to reduced vesicular markers at the fusion focus and formation of ectopic foci. **A-E.** Myo52-tdTomato and (A) Exo84-GFP, (B) Exo70-GFP, (C) GFP-Ypt3, (D) Agn2-sfGFP and (E) Eng2-sfGFP in WT and *acp2Δ*, before fusion time. White arrow highlight ectopic foci. Note that these images, selected to show ectopic foci, stem from various timepoints during time-lapse imaging, for which specific timing in the fusion process and photobleaching may mask the difference in fusion focus intensity. **F.** Boxplot of fusion focus fluorescence intensities at fusion time normalized to WT in the strains mentioned in (A-E). p-values relative to WT are 3.7×10^−11^, 4.2×10^−7^, 3.4×10^−8^, 1.2×10^−9^, 2.5×10^−5^ and 1.9×10^−1^ for Myo52, Exo84, Exo70, Ypt3, Agn2 and Eng2 in *acp2*Δ, respectively (for both strains, n = 40, 25, 40, 30, 11 and 11 cells, respectively). **G.** Time-lapse images of Myo52-tdTomato in WT and *acp2*Δ during the fusion process. White arrows show ectopic Myo52 foci. Time in hour:min. **H**. Boxplot quantifying the number of time frames at which a Myo52 ectopic focus was observed during the fusion process in time-lapse imaging as in (G). p-values relative to WT are 4.8×10^−3^, 8.7×10^−8^ and 7.0×10^−5^ for *acp1Δ, acp2Δ* and *acp1Δacp2Δ*, respectively. This was measured on 32, 53, 88 and 56 individual mating pairs for WT, *acp1Δ, acp2Δ* and *acp1Δacp2Δ*, respectively. Bars are 5µm.

The question is why the more actin-rich *acp2*Δ fusion foci accumulate fewer exocytic vesicles. Looking closely at Myo52, we noticed that *acp2Δ* cells frequently form Myo52 foci away from the fusion focus (Figure 3G). Such ectopic Myo52 foci formed in both P- and M-cells, repeatedly during the fusion process. In time-lapse imaging of the fusion process at 5 min intervals, *acp2Δ* cells displayed on average 7 time points with ectopic foci, *acp1Δ* and *acp1Δ acp2Δ* cells displayed 1, and we barely found any for WT cells with this set-up (Figure 3H). Note that upon camera upgrade (which happened in the course of the project) or when using spinning disk imaging, more ectopic foci became detectable in all backgrounds, including WT. The detection of rare, transient ectopic foci in WT mating pairs (see Figure 1C, 10 min time-frame, for example) suggests that the behaviour exhibited by *acp2Δ* cells occurs but is normally repressed in WT cells. These ectopic Myo52 foci extensively colocalized with the exocyst subunits Exo84 and Exo70, and the Rab11 GTPase Ypt3 (Figures 3A-C, Figure S3G-I). We could not detect glucanases at Myo52 ectopic foci, likely because glucanase sfGFP tagging partly impairs their function and their expression levels are very low (Figure 3D-E). Put together, these results suggest that the formation of ectopic foci is the likely cause of the reduced amounts of Myo52 and cargoes at the fusion focus.

**Figure S3 – related to Figure 3.**
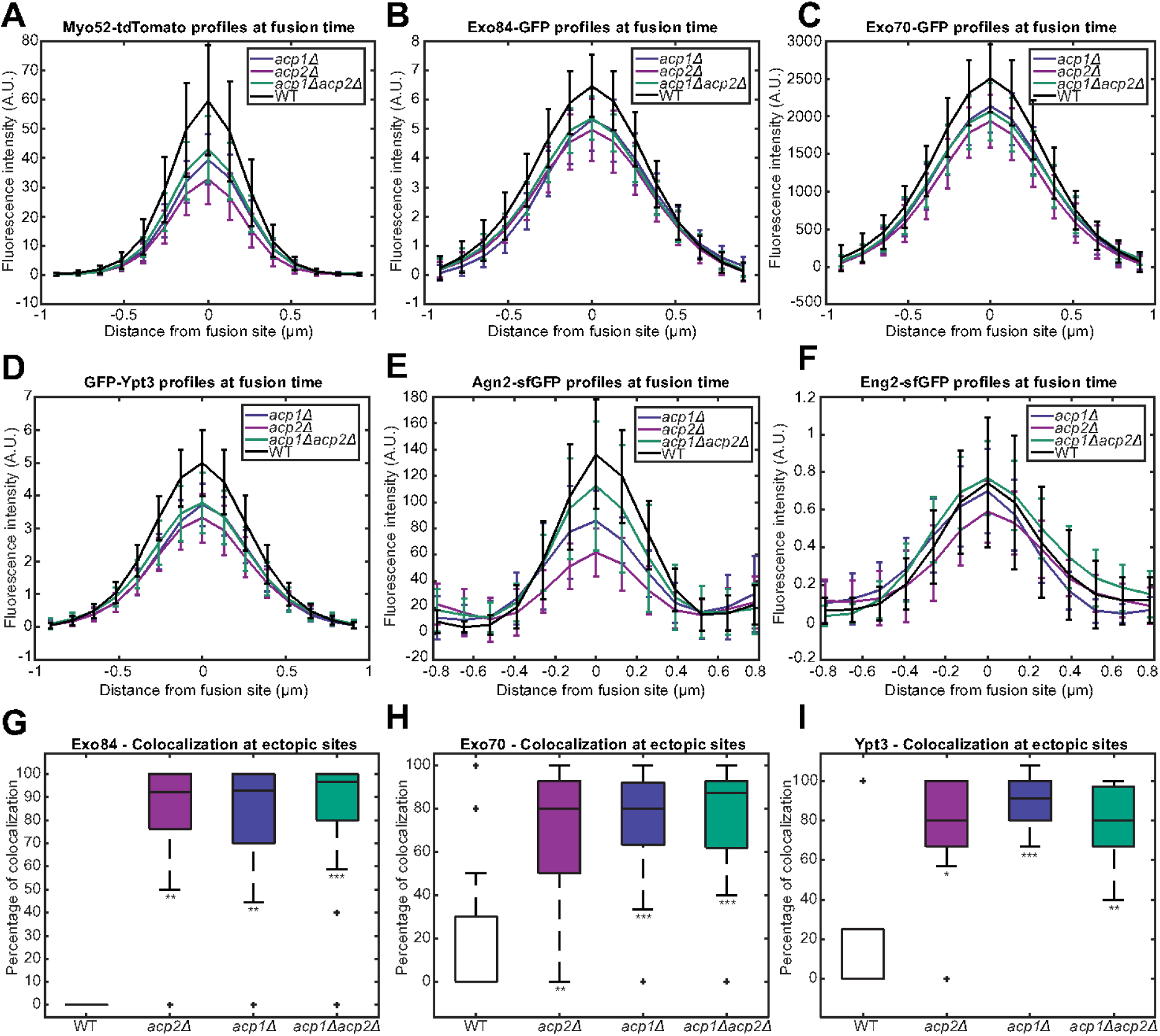
Quantifications of vesicular marker intensities at the fusion focus and ectopic foci in absence of CP. **A-F.** Profiles of the bleach-corrected fluorescence intensities around the fusion focus at fusion time in the strains shown in Figure 3. The boxplot in Figure 3F shows the central points of these profiles further normalized to WT. (A) Myo52-tdTomato. (B) Exo84-GFP (C) Exo70-GFP. (D) GFP-Ypt3. (E) Agn2-sfGFP. (F) Eng2-sfGFP. **G-I.** Boxplots of the colocalization of Myo52 with (G) Exo84, (H) Exo70, and (I) Ypt3 at ectopic sites in WT, *acp1Δ, acp2Δ*, and *acp1Δ acp2Δ* strains. p-values relative to WT are 1.1×10^−7^, 8.7×10^−7^ and 6.0×10^−9^ for Exo84, 1.3×10^−5^, 2.9×10^−9^ and 3.5×10^−9^ for Exo70 and 3.1×10^−7^, 8.4×10^−9^ and 3.5×10^−6^ for Ypt3, for *acp1Δ, acp2Δ* and *acp1Δacp2Δ*, respectively. This was measured on 7, 16, 20, and 20 individual cells for WT, *acp1Δ, acp2Δ* and *acp1Δacp2Δ* for Exo84, 28 individual mating pairs for each strain for Exo70 and 9, 29, 28, and 27 individual mating pairs for WT, *acp1Δ, acp2Δ* and *acp1Δacp2Δ*, for Ypt3.

### Myo52 ectopic foci form at actin patches

Using higher speed time lapse spinning disk imaging at 1 s intervals, we found that Myo52 ectopic foci did not appear randomly: they always formed at the cell periphery, rarely moving away from it; they did not break-off from the fusion focus; instead they formed at remote locations, occasionally moving back and fusing with the fusion focus (Figure 4A; Movie S1). The colocalization with exocytic markers stood true at this higher temporal resolution, as exemplified by Exo70 (Figure 4B). These observations suggest that ectopic foci are nucleated at remote location in absence of CP.

**Figure 4.**
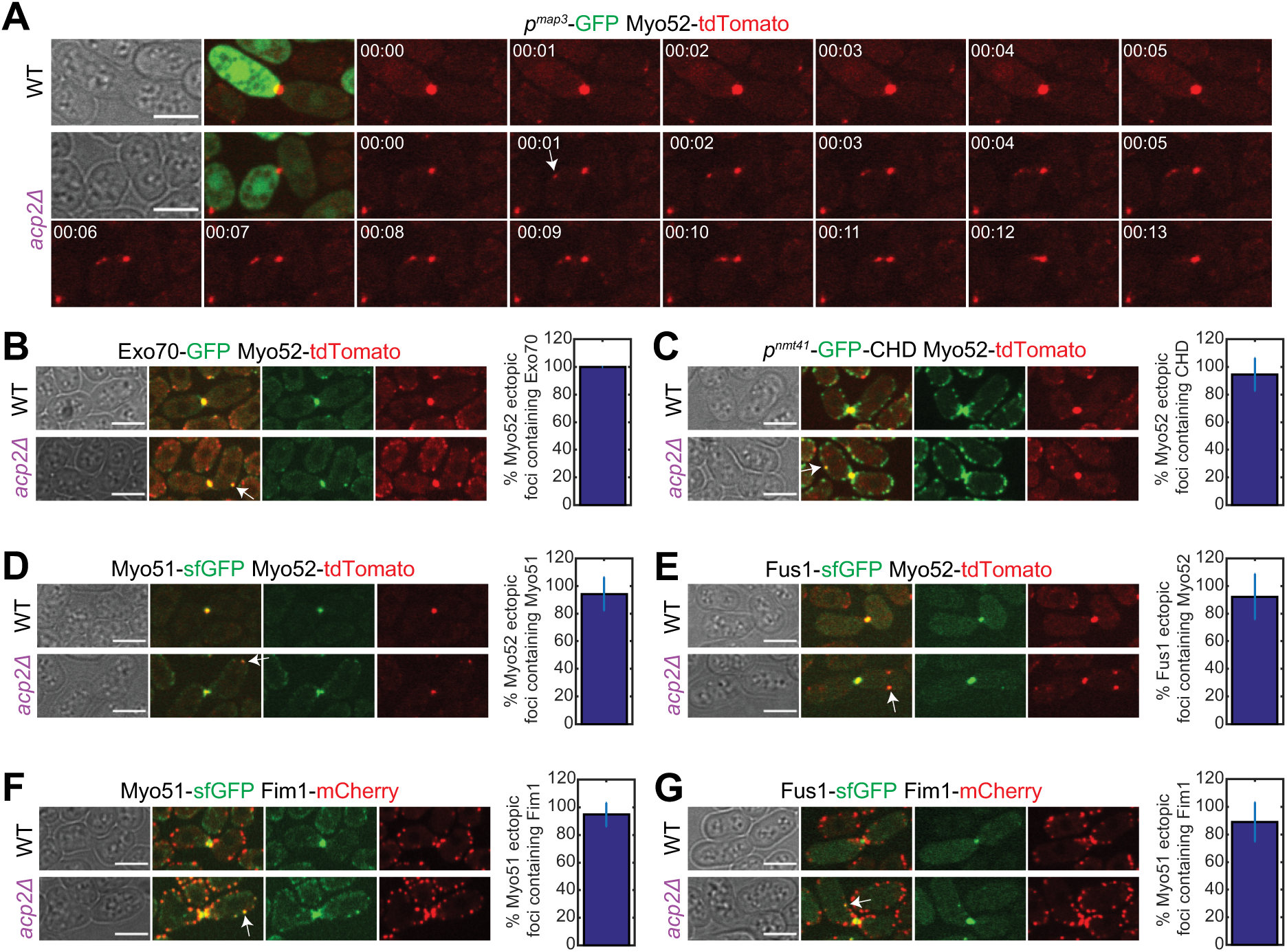
Myo52 ectopic foci form at actin patches. **A.** Spinning-disk confocal microscopy time-lapse images of Myo52-tdTomato and cytosolic GFP expressed under the *map3* promoter (shown only for first time point) in WT and *acp2Δ* before fusion time. The white arrow marks an ectopic Myo52 focus that forms at 1s and then moves towards the fusion focus. Time in min:sec. **B-E.** Spinning-disk confocal microscopy images of Myo52-tdTomato and (B) Exo70-GFP, (C) GFP-CHD, (D) Myo51-sfGFP, and (E) Fus1-sfGFP in WT and *acp2Δ* before fusion time. **F-G.** Spinning-disk confocal microscopy images of strains expressing Fim1-mCherry and (F) Myo51-sfGFP or (G) Fus1-sfGFP in WT and *acp2Δ* before fusion time. White arrows mark ectopic Myo52, Fus1 and Myo51 foci. The bar plot to the right of the images shows the proportion of ectopic foci colocalizing with indicated markers, of which an example is shown with a white arrow. Bars are 5µm.

Interestingly, using GFP-CHD as F-actin marker, we found that more than 90 % of the ectopic Myo52 foci colocalized, at least transiently, with what looked like actin patches (Figure 4C). Ectopic Myo52 foci also colocalized with Myo51, a second type V myosin that normally associates with Rng8, Rng9 and tropomyosin to decorate linear actin structures and plays a structural role in the coalescence of the focus [44, 51, 52] (Figure 4D). Whereas Myo51 principally decorates the fusion focus in WT cells, in *acp2*Δ it additionally prominently localized to actin patches marked by the specific actin patch marker Fim1, which bundles actin filaments [53, 54] (Figure 4F). Remarkably, the better signal-to-noise ratio of spinning disk microscopy also revealed the presence of Fus1 in Myo52 ectopic foci (Figure 4E; Movie S2), which colocalized with Fim1 (Figure 4G; Movie S3). Thus, the absence of CP leads to the formation of ectopic foci of formin Fus1 and type V myosin at Arp2/3-nucleated actin patches.

The presence of Fus1 at actin patches suggests that exposed filament barbed ends in absence of CP are bound by Fus1, which nucleates ectopic foci, leading to the observed *acp2Δ* phenotypes. To investigate whether CP’s capping function is required to protect patches from Fus1, we deleted Acp1 and Acp2 tentacles (*acp1*^*Δt*^ and *acp2* ^*Δt*^) to reduce CP affinity for actin barbed ends. These truncations compromised localization to actin patches, with Acp2^Δt^ retaining more actin patch binding than Acp1^Δt^ (Figures 5B, E, H), in agreement with previous in vitro work [14]. They also showed more frequent ectopic Myo52 foci than WT cells, in an order consistent with their retained actin binding capacity (Figure 5B, E, I). To probe whether CP localization to actin patches is required to protect against Fus1, we generated *acp2*^*R12A,Y77A*^, an allele predicted not to bind the CPI motif present on most CP interactors [18]. This allele localized inefficiently to actin patches (Figure 5G, H), and exhibited ectopic foci and increased fusion times, to levels intermediary between WT and *acp2Δ* cells (Figure 5J). These results indicate that CP needs to bind F-actin in patches to prevent the formation of ectopic foci.

**Figure 5.**
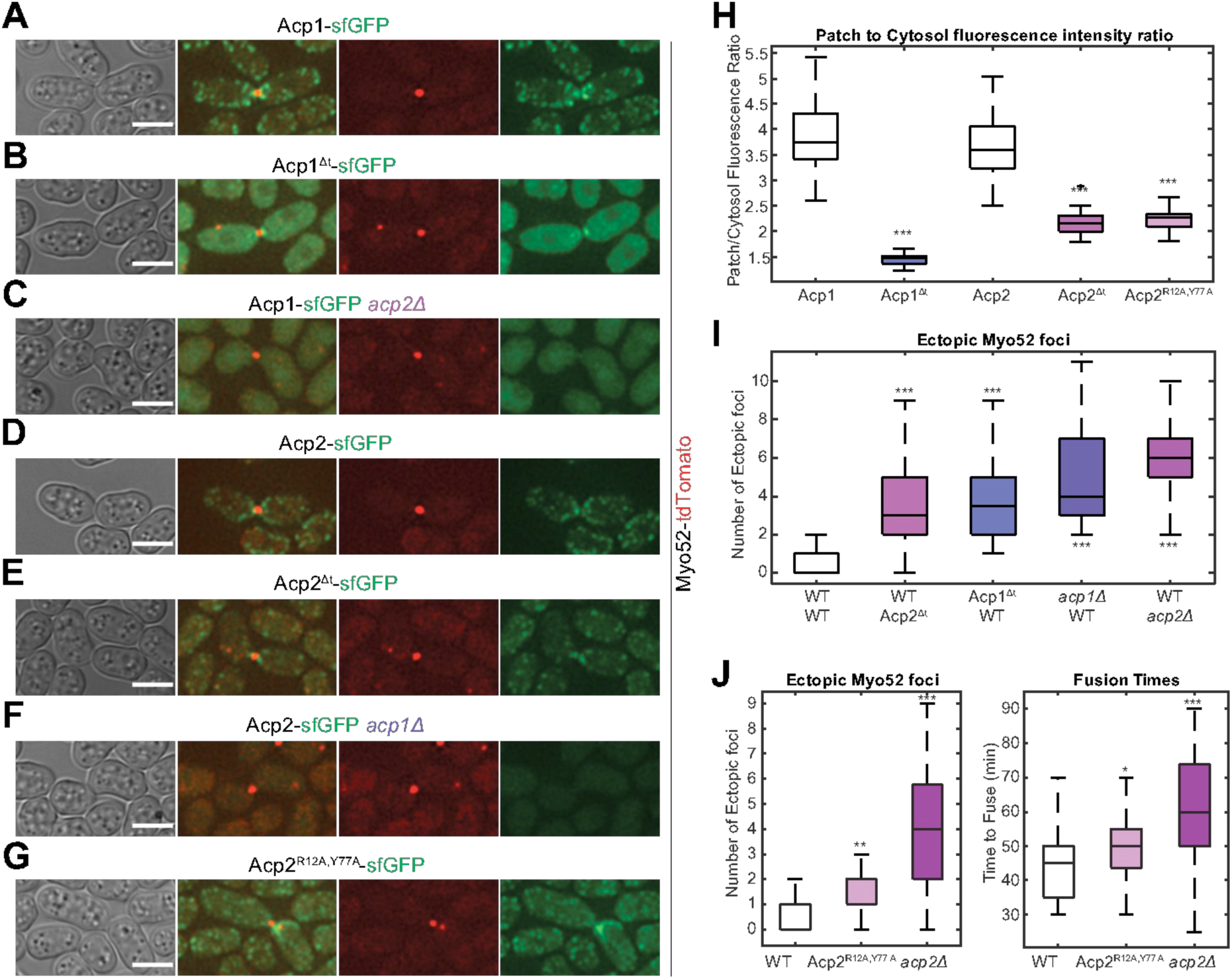
CP recruitment to actin patches and barbed end binding are required to protect against ectopic foci. **A-G.** Myo52-tdTomato and sfGFP-tagged capping protein subunit (Acp1 and Acp2), lacking or not its tentacle (Δt), carrying mutated CPI-interacting residues (R12A Y77A), or in absence of the other subunit at fusion time, as indicated. Bars are 5µm. **H.** Boxplot of Acp1-sfGFP or Acp2-sfGFP patch to cytosol fluorescence intensity ratios in fusing cells of strains as in (A-G). p-values relative to the corresponding WT are 6.0×10^−34^, 1.8×10^−20^ and 1.3×10^−21^ for Acp1^*Δt*^-sfGFP, Acp2^*Δt*^-sfGFP and Acp2^*R12A,Y77A*^-sfGFP, respectively (n = 36 individual mating pairs for each strain). **I.** Boxplot quantifying the number of time frames at which a Myo52 ectopic focus was observed during the fusion process in time-lapse imaging of strains as in (A-F). p-values relative to WT are 3.0×10^−18^, 4.2×10^−24^, 4.6×10^−33^ and 1.3×10^−45^ for Acp2^*Δt*^-sfGFP, Acp1^*Δt*^-sfGFP, *acp1Δ* and *acp2Δ* respectively (n = 34 individual mating pairs for each strain). **J.** Boxplots of ectopic foci (as in I) and fusion times for WT (*acp2-sfGFP*), *acp2*^*R12A,Y77A*^*-sfGFP* and *acp2Δ* strains. p-values relative to WT are 2.2×10^−5^ and 3.2×10^−18^ for ectopic foci for *acp2*^*R12A,Y77A*^*-sfGFP* and *acp2Δ* respectively, and 2.5 10^−4^ and 4.7 10^−13^ for fusion times. This was measured on 65, 57 and 59 individual mating pairs for WT, Acp2^R12A,Y77A^-sfGFP and *acp2Δ* respectively.

Finally, to test whether ectopic foci cause the fusion delay by diverting vesicles away from the cell-cell contact zone, we artificially recruited Myo52-GFP to patches labelled with Fim1-GBP-mCherry by using the high affinity between GFP-Binding Protein (GBP) and GFP (Figure S4). This led to fusion delay and persistence of the fusion focus after fusion, replicating the *acp2Δ* phenotypes (Figure 3H, Figure S4).

Together, these results show that during cell fusion, CP insulate actin patches from Fus1. This ensures that Fus1 activity is restricted to, and all myosin V-driven cargoes directed to, the site of cell-cell contact.

### Uncapped actin patches recruit formins and acquire dual identity in interphase cells

To test whether CP more generally protects actin patch identity, we investigated the influence of CP deletion on actin-binding proteins in interphase cells. Previous work had shown that actin patches are dispersed in the cell in absence of CP both in *S. cerevisiae* [55] and in *S. pombe* [40, 56]. *S. pombe* cells lacking CP also exhibit weak actin cables [33, 40]. Remarkably, markers normally associated with formin-nucleated actin cables were perturbed in absence of CP: While Myo51 labels cable-like structures in WT interphase cells [52], it formed punctate structures that colocalized with Fim1 at both cell tips and sides in *acp2Δ* cells (Figure 6A). Similarly, Myo52, which mainly localizes to cell tips in WT cells, formed dots on the tips and sides of *acp2Δ* cells, which coincided with Myo51 and Fim1 patches (Figure 6B-C). Finally, tropomyosin Cdc8, which stabilizes actin cables and is largely absent from actin patches in WT cells, formed foci that colocalized with Fim1 in *acp2Δ* cells (Figure 6D; Movie S4). The co-incidence of fimbrin and tropomyosin is particularly remarkable given recent data showing that competition between these two proteins drives their sorting to distinct Arp2/3 and formin-nucleated networks, respectively [9]. Thus, actin patches assume a dual identity in absence of CP both during mating and during vegetative growth.

**Figure 6.**
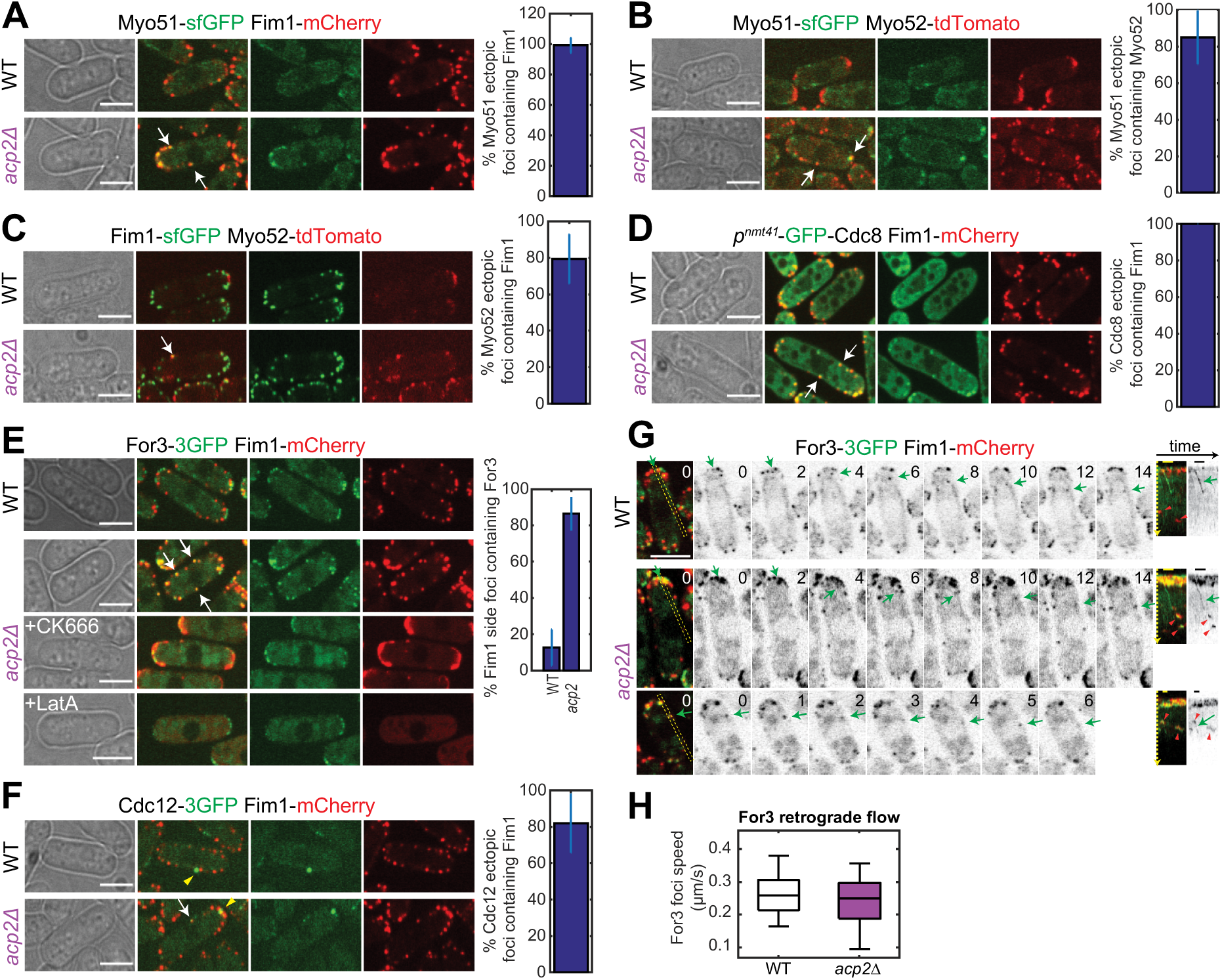
Capping proteins insulate actin patches from Myo52 and actin cable markers in interphase cells. **A-F.** Spinning-disk confocal microscopy images of (A) Fim1-mCherry and Myo51-sfGFP, (B) Myo52-tdTomato and Myo51-sfGFP, (C) Myo52-tdTomato and Fim1-sfGFP, (D) Fim1-mCherry and GFP-Cdc8, (E) Fim1-mCherry and For3-3GFP, and (F) Fim1-mCherry and Cdc12-3GFP, in WT and *acp2*Δ interphase cells during exponential growth. In (E), the bottom panels show *acp2*Δ cells treated with 500 µM CK-666 for 5 min or 200µM Latrunculin A for 5 min. White arrows highlight colocalization events in *acp2*Δ, which do not occur in WT cells. Yellow arrowheads point to Cdc12 spots. The proportion of colocalization at ectopic sites along the cell sides is shown with the bar plot to the right to the images. **G.** Spinning-disk confocal time-lapse images of Fim1-mCherry and For3-3GFP (green and grey) showing retrograde flow in WT (top) and *acp2*Δ cells (bottom two panels). The bottom panel shows an example of retrograde flow starting at a lateral actin patch. On the right of the timelapse are kymographs of the yellow dashed boxed region. Green arrows point to For3-3GFP retrograde flow movement. Red arrowheads in kymographs show lateral actin patches on which For3 localizes in *acp2*Δ but not WT cells. **H.** Boxplot showing the For3 retrograde flow rate. p-value is 0.81 (n=35 For3 retrograde movements in 68 cells for WT and 17 in 340 cells for *acp2*Δ imaged over 3 min). Time in seconds. Bars are 5µm.

**Figure S4 – related to Figure 5.**
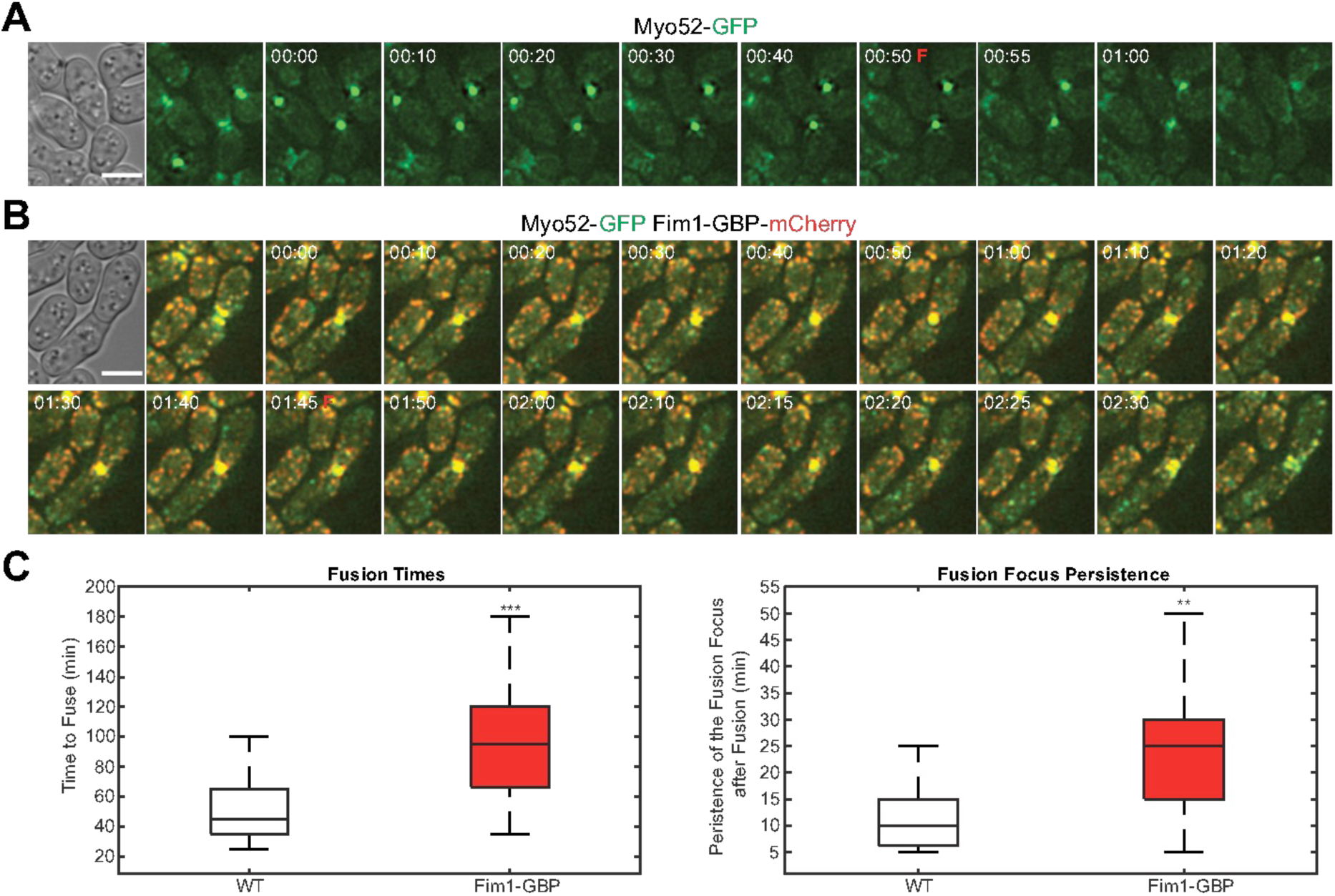
Forced recruitment of Myo52 to actin patches delays fusion. **A-B.** Time-lapse images of a strain expressing Myo52-GFP alone (A) or in combination with Fim1-GBP-mCherry (B) from beginning to disappearance of the fusion focus. The beginning is defined as the first formation of the focus in both cells. The fusion point is highlighted next to the time with a red F and is defined as the peak Myo52-GFP fluorescence intensity at the focus. Time in hour:min. Bars are 5µm. **C.** Boxplots of the above-mentioned strain fusion times and focus persistence times. p-values relative to WT are 2.1×10^−8^ and 9.5×10^−8^ for fusion times and persistence times, respectively (n = 55 mating pairs for each strain).

Because Fus1 is not expressed during mitotic growth [42], we monitored the localization of For3 and Cdc12 formins. For3, which only occasionally overlapped with actin patches in WT cells, was prominently present at actin patches all over the cell in *acp2*Δ mutants (Figure 6E; Movie S5). Disruption of patches by treatment with the Arp2/3 inhibitor CK-666 or full actin depolymerization with Latrunculin A restored For3 localization to cell poles (Figure 6E). Arp2/3 inhibition also led to Fim1 relocalization to cell poles. This re-localization is likely due to Fim1 recruitment to excess formin-assembled cables in absence of actin patches [3], as complete actin depolymerization rendered Fim1 cytosolic (Figure 6E). Thus, uncapped actin patches ectopically recruit For3.

Cdc12 also formed ectopic foci that coincided with actin patches in *acp2*Δ cells (Figure 6F). Note that these ectopic foci were distinct from the previously reported spot of Cdc12 [57], which occurs also in WT cells, where it does not coincide with patches, has a longer life-time, and is more intense (Figure 6F). To address whether formins are active at patches, we first tested whether their inactivation would alleviate the localization of tropomyosin at *acp2*Δ actin patches. However, deletion of *for3* by itself led to significant Cdc8 enrichment on actin patches, likely because F-actin and associated protein homeostasis is perturbed in absence of actin cables (Figure S5A). Therefore, not surprisingly, Cdc8 also localized to actin patches in *for3*Δ *acp2*Δ and *cdc12-112 for3*Δ *acp2*Δ mutants (Figure S5). We then directly probed for For3 activity by observing its retrograde flow, which depends on the assembly of actin in cables [46]. For3 retrograde flow occurred at similar rates in WT and *acp2*Δ cells, though fewer movements were observed in *acp2*Δ cells, consistent with the weak actin cables in these cells (Figure 6G-H) [33, 40]. Interestingly, For3 retrograde movements could be observed to initiate from actin patches, indicating assembly of cables from the patches (Figure 6G). We conclude that, independently of the specific formin, CP insulates actin patches from formins, restricting their activity to the proper location.

**Figure S5 – related to Figure 6.**
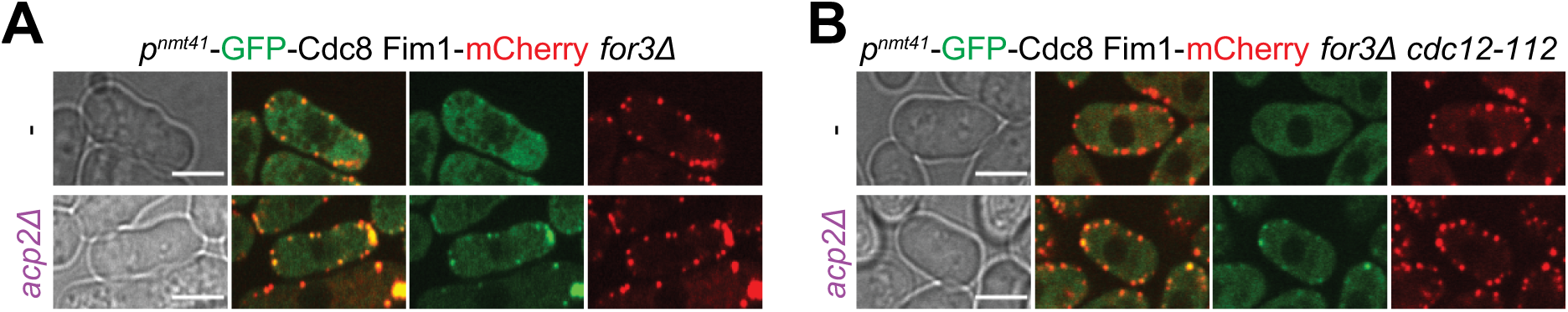
Removing formins does not lead to the recovery of actin patch identity. **A-B.** Spinning-disk confocal microscopy images of Fim1-mCherry and GFP-Cdc8 in interphase cells during exponential growth in (A) *for3Δ* and (B) *for3Δ cdc12-112* at 36°C, in otherwise WT and *acp2*Δ background. Bars are 5µm.

## Discussion

How cells simultaneously assemble functionally diverse actin structures of distinctive identity within a common cytosol is a debated question. The Arp2/3 complex and formins respectively assemble branched and linear actin structures, which are decorated by largely distinct sets of actin-binding proteins. A hallmark of formins is their ability to promote barbed end growth against the growth-arrest function of capping protein (CP), a feature that has been demonstrated in numerous in vitro studies [27–32, 58–60]. However, how this competition plays out in vivo, and how CP may prevail against formins was largely unexplored. In this work, we have shown that CP actively protects Arp2/3-assembled actin patches against formins, thus preserving their identity and restricting formin to their proper location.

Our interest in CP arose from the finding that CP mutants exhibit a delay in cell-cell fusion. For fusion, cells locally digest their cell wall by concentrating the delivery of secretory vesicles carrying cell wall hydrolases at the site of cell-cell contact [41]. The actin fusion focus assembled by the formin Fus1 concentrates secretory vesicles marked by type V myosin Myo52, Ypt3/Rab11 GTPase and the exocyst complex. By contrast, in cells lacking CP, despite strong accumulation of Fus1 at the fusion site, secretory vesicles are frequently diverted away from the fusion site to ectopic Fus1 foci at actin patches. This correlates with a reduction in secretory vesicle markers and their hydrolase cargoes at the fusion focus, which likely explains the observed delay in fusion. Consistent with this view, the forced diversion of Myo52 to actin patches also yields extended fusion times. Thus, the fusion defect of cells lacking CP likely results from the diversion of secretory vesicles to ectopic foci-like structures, rather than from excessive actin assembly at the fusion site.

This raises the question of where CP acts – in formin-assembled structures or at Arp2/3-assembled actin patches. We argue that competition between formins and CP happens throughout the cell, with distinct outcome: in formin-assembled structures, the formin wins and CP is largely dispensable; at actin patches, CP dominates and is essential to insulate this structure against all formins. These distinct outcomes are reflected in the exclusive localization of CP at patches and restriction of formins to other cellular locations. The occasional ectopic foci detected in WT cells illustrate that CP patch protection against formins is an active process that can transiently fail even in WT cells.

CP-formin competition is best revealed in cells in which it is compromised by alteration of Fus1 and/or CP function. At the fusion focus, the competition is best revealed by the examination of Fus1 FH2 mutations. A first interesting observation is that these mutations compromised fusion to various extent in vivo, although they all fully abrogated actin assembly in vitro [50]. Thus, the in vitro activity is poorly predictive of the specific activity in vivo, likely because the local Fus1 concentration in the focus is vastly exceeds the concentrations tested in vitro. A second important observation is the strong allele-specific suppression by *acp2*Δ of the fusion defect conferred by *fus1*^*I951A*^, which carries a mutation in the knob region. At the molecular level, this allele-specific suppression is consistent with the proposed structural arrangement of the formin-CP ternary complex at the barbed end, which shows a steric clash between the formin knob and CP beta-subunit [31]. By blocking the association of the formin knob with the barbed end, the I951A mutation likely favors the binding of the CP beta-subunit, thus promoting capping function in the ternary complex. At the cellular level, the inability of Fus1^I951A^ to assemble a fusion focus must be due to CP successfully competing with the formin at the fusion site. Further evidence for competition at this location comes from the increased intensity of Fus1 at the fusion focus and the extended lifetime of the fusion focus in CP mutant cells. This suggests that small amounts (below detection levels) of CP may compete with Fus1 in the fusion focus, although the increased Fus1 intensity may also be due to Fus1 recruitment to adjacent actin patches, which closely associate with the focus. We conclude that some Fus1-CP competition can take place at the fusion site, although Fus1 normally dominates, whether CP is present or not.

Indeed, our data indicate that the principal sites of formin-CP competition are actin patches, where CP efficiently outcompetes Fus1. In WT cells, CP is strongly enriched at actin patches and prevents formin binding. In cells lacking CP function, all three formins localize to actin patches. The strength of CP protection against formins largely scales with its barbed end binding affinity as measured in vitro [14], with *acp2*^Δ*t*^ showing fewer ectopic foci than *acp1*^Δ*t*^, and *acp1*Δ or *acp2*Δ. When CP is absent from patches, these acquire characteristics normally specific to linear actin structures: they are decorated by myosin V Myo51 and tropomyosin, which normally preferentially associates with actin cables, ring and focus [44, 51, 52] [7, 43, 47–49]; they also accumulate the myosin V Myo52, which erroneously transports its cargoes to these locations. The coincidence of tropomyosin and fimbrin at patches devoid of CP is particularly striking given the previous findings that competition between these two actin-binding proteins is sufficient for their specific association with formin- and Arp2/3-assembled structure in vitro [9–11]. We conclude that, in absence of CP, actin patches acquire a double identity.

One important question is whether the double identity of CP-devoid patches arises from ectopic local formin activity or simply from uncapping. This question is difficult to address given that any perturbation in actin structures will perturb homeostasis [3]. For instance, deletion of formins frees G-actin and tropomyosin that can now incorporate in actin patches. Conversely, disruption of actin patches enhances formin-assembled structures, which become more permissive for fimbrin association. However, two observations argue for activity of formins at patches lacking CP. First, during mating, Myo52 was not present at all actin patches, but was always there when Fus1 was. This argues that Fus1 activity is the driving force for the recruitment of vesicular cargoes to patches. Second, the For3 retrograde flow from patches indicates assembly of cables from this location [46]. As a side note, while the mechanism and function of For3 retrograde flow remain unknown, the observation of For3 retrograde flow at similar rate in cells lacking CP indicates that it is not due to arrest of For3-dependent barbed end growth upon formation of a ternary formin-CP complex. In summary, observations indicate that the now patch-localized formins are active in assembling linear actin filaments.

An interesting observation is that *acp2*Δ cells consistently displayed stronger phenotypes not only than *acp1*Δ, but also than *acp1*Δ *acp2*Δ double mutants. Berro and Pollard also previously noted that the phenotypes of *acp2*Δ and *acp1*Δ are not identical [24]. In light of the protective role of CP against formins, one interpretation is that, while formins gain access to actin barbed ends in absence of Acp2, they may gain better access if Acp1 is still present. Because Acp1 still weakly binds the actin barbed end in absence of Acp2 in vitro [16], its presence may somehow help recruit formin to the barbed end, causing the more pronounced phenotype. This interpretation predicts an interaction interface between formins and the CP alpha subunit, a hypothesis consistent with the recently described association of human INF2 formin with the CP alpha subunit [61].

Because CP-formin competition yields distinct outcome at the sites of formin action and Arp2/3-assembled patches, one important question is what defines the competition outcome. Part of the answer comes from our investigation of an Acp2 allele carrying mutations in the CPI-binding residues (R12A,Y77A), which compromises CP localization to actin patches. This finding agrees with previous data in human cells that interaction of CP with CPI-containing proteins contributes to CP localization. The specific CP binding partners in actin patches are unknown in *S. pombe*, but may involve the homologue of *S. cerevisiae* Aim21, which was recently proposed to bind CP through the CPI-binding residues and contribute to its localization to actin patches [19], though this finding was not reproduced in a second study [20]. Consistent with this hypothesis, deletion of *S. pombe* Aim21 was identified in our genome-wide screen to have fusion defects [44]. In any case, the phenotype of *acp2*^*R12A,Y77A*^ cells indicate that CP recruitment to actin patches by pre-localized binding partners is required to protect them against formin activity. Thus, barbed end-independent recruitment of CP may tip the CP-formin competition in favour of CP in Arp2/3-assembled structures.

The findings described in our study present the CP-formin competition in a new light, where the principal role of CP is to protect Arp2/3 structures against ectopic localization of formins. In fission yeast *acp2Δ* cells, the inappropriate localization of formins to actin patches has important consequences on cellular organization: during mating, Fus1 diverts cargoes away from the fusion site, slowing the fusion process; in interphase cells, For3 localization to patches is the likely cause of the previously noted actin cables disorganization and partial loss of cell polarity [24, 33]; in dividing cells, the reason for the cytokinetic defect of cells lacking CP [33] may also be the titration of Cdc12 to actin patches. As CP is ubiquitous in eukaryotic cells, it likely protects the identity of Arp2/3 actin assemblies and prevents formin ectopic activity in a vast range of organisms.

## Materials and Methods

### Contact for reagent and resurce sharing

Further information and requests for resources and reagents should be directed to and will be fulfilled by the Lead Contact, Sophie Martin.

### Experimental model and subject details

*S. pombe* strains used in this study are listed in Table S1. Homothallic (h90) strains able to switch mating types were used. For mating experiments, cells were grown in liquid or agar Minimum Sporulation Media (MSL), with or without nitrogen (+/−N) [62, 63]. For interphase experiments, cells were grown in liquid or agar Edinburgh minimal medium (EMM) supplemented with amino acids as required.

### Method details

#### STRAIN CONSTRUCTION

Strains were constructed using standard genetic manipulation of *S. pombe* either by tetrad dissection or transformation.

Genes were tagged at their endogenous genomic locus at their 3’ end, yielding C-terminally tagged proteins, or deleted by replacing their ORF by a resistance cassette. This was achieved by PCR amplification of a fragment from a template plasmid with primers carrying 5’ extensions corresponding to the last 78 coding nucleotides of the ORF (for C-terminal tagging) or the last 78 nucleotides of the 5’UTR (for gene deletion) and the first 78 nucleotides of the 3’UTR, which was transformed and integrated in the genome by homologous recombination, as previously described [64]. For tagging of genes with sfGFP, a pFA6a-sfGFP-kanMX plasmid (pSM1538) was used as a template for PCR-based targeted tagging of *myo51, fim1, acp1* and *acp2*. For tagging of genes with mCherry, a pFA6a-mCherry-natMX plasmid (pSM684) was used as a template for PCR-based targeted tagging of *fim1*. For tagging of genes with GBP-mCherry, a pFA6a-GBP-mCherry-natMX plasmid (pSM1768) was used as a template for PCR-based targeted tagging of *fim1*. For PCR-based targeted deletion, a pFA6a-hphMX plasmid (pSM693) or a pFA6a-bleMX plasmid (pSM694) was used as a template for *acp1* and *acp2* respectively. For deletion of the tentacle of Acp1 and Acp2, PCR-based targeted tagging was done using the last 78 nucleotides upstream of the C-terminal tentacle (R233-T256 for Acp1 and R244-I268 for Acp2).

For point mutations of *acp2* (R12A,Y77A) and *fus1* (K879A, I951A, GN1087,1088RP, K1112A), site directed mutagenesis was conducted on plasmids constructed as followed 5’UTR-ORF-sfGFP-3’UTR by PCR amplification of the full fragments from strains IBC180 or IBC135 with primers carrying unique restriction sites, which was then cloned into a pSP72 plasmid (pSM1232). After site-directed mutagenesis, the resulting plasmids (pSM2203, pSM2251, pSM2252, pSM2253 and pSM2254, respectively) were sequenced, linearized and transformed into recipient strains YSM2440 for *acp2* and IBC178 (*fus1Δ* strain) for *fus1* mutants.

Construction of the strain overexpressing *fus1* (p^nmt1^-fus1-sfGFP) was done by integration of fus1-sfGFP under the *nmt1* promoter at the *ura4+* locus. First, the *nmt1* promotor was amplified from a pREP1 plasmid (pSM1758) with primers carrying KpnI and NotI extensions. Second, the fus1-sfGFP fragment was amplified from strain IBC180 with primers carrying NotI and SacI extensions. These two fragments were cloned by 3-point ligation into the vector pAV133 (pJK211, a kind gift from Dr. Aleksandar Vjestica, UNIL) digested with KpnI and SacI. The resulting plasmid (pSM2282) was sequenced, digested with AfeI and stably integrated as a single copy at the *ura4+* locus into strain YSM2440.

Construction of the strain expressing ypt3 (p^nmt41^-GFP-ypt3) was done by integration of GFP-ypt3 under the *nmt41* promoter at the *ura4+* locus. The p^nmt41^-GFP-ypt3 fragment was digested from a pREP41-GFP-ypt3 plasmid (pSM893) with PstI and XmaI and cloned into the vector pAV133 (pJK211, a kind gift from Dr. Aleksandar Vjestica, UNIL) digested with the same enzymes. The resulting plasmid (pSM2250) was linearized by AfeI and transformed into strain YSM2440.

#### MATING ASSAYS

Live imaging of *S. pombe* mating cells protocol was adapted from [63]. Briefly, cells were first pre-cultured overnight in MSL+N at 25°C, then diluted to OD_600_ = 0.05 into MSL+N at 25°C for 20 hours. Exponentially growing cells were then pelleted, washed in MSL-N by 3 rounds of centrifugation, and resuspended in MSL-N to an OD_600_ of 1.5. Cells were then grown 3 hours at 30°C to allow mating in liquid, added on 2% agarose MSL-N pads, and sealed with VALAP. We allowed the pads to rest for 30 min at 30°C before overnight imaging, or for 3 h before high speed imaging.

For interphase imaging, cells were grown to exponential phase at 30°C in EMM+ALU media, pelleted and added to 2% agarose EMM+ALU pads.

For CK-666 (Sigma) and LatA (Enzo Life Sciences) treatment, the drugs were added directly before imaging to the final resuspension, to a final concentration of 500µM and 200µM, respectively. The slide was allowed to rest for 5 minutes before imaging.

#### MICROSCOPY

Images presented in Figures 1, 2, 3, 5, and S2, S4, except panel 2E were obtained using a DeltaVision platform (Applied Precision) composed of a customized inverted microscope (IX-71; Olympus), a UPlan Apochromat 100×/1.4 NA oil objective, a camera (CoolSNAP HQ2; Photometrics or 4.2Mpx PrimeBSI sCMOS camera; Photometrics), and a color combined unit illuminator (Insight SSI 7; Social Science Insights).

Figures were acquired using softWoRx v4.1.2 software (Applied Precision). Images were acquired every 5 minutes during 9 to 15 hours. To limit photobleaching, overnight videos were captured by optical axis integration (OAI) imaging of a 4.6-µm z-section, which is essentially a real-time z-sweep.

Images presented in Figures 4, 6, and S5 and panel 2E were obtained using a spinning-disk microscope composed of an inverted microscope (DMI4000B; Leica) equipped with an HCX Plan Apochromat 100×/1.46 NA oil objective and an UltraVIEW system (PerkinElmer; including a real-time confocal scanning head [CSU22; Yokagawa Electric Corporation], solid-state laser lines, and an electron-multiplying charge coupled device camera [C9100; Hamamatsu Photonics]). Time-lapse images were acquired at 1s interval using the Volocity software (PerkinElmer).

### Quantification and statistical analysis

Fusion efficiencies were calculated as in [41]. Briefly, at the specified time post-starvation, mating pairs and fused pairs were quantified using the ImageJ Plugin ObjectJ, and the subsequent fusion efficiency was calculated using the following equation:

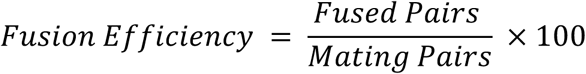

Fusion Times were calculated using the 5-minutes time lapse overnight movies using the 2-dots Myo52-tdTomato stage [41] as a marker for the beginning of the fusion process and either the entry of GFP expressed under control of the P-cell-specific *p*^*map3*^ promoter into the h-partner, or the maximum intensity of the Myo52-tdTomato dot, the two of which perfectly correlate [41], as a marker for the end of the process. Persistence times were calculated using the 5-minutes time lapse overnight movies using fusion time as beginning and last appearance of the Myo52-tdTomato dot as end of the post-fusion focus lifetime. Fusion times and persistence times vary between experiments because, for all experiments but panels 1A, C, only early timepoints were considered to avoid Myo52 bleaching, which most certainly induces a bias toward quickly fusing cells. The WT control was imaged and quantified for each experiment to allow comparison within an experiment.

Fusion Focus intensities at fusion time were obtained using the 5-minutes time lapse overnight movies using either the entry of GFP into the h-partner, or the maximum intensity of the Myo52-tdTomato dot to determine the moment of fusion. On that time frame, a fluorescence profile across the fusion focus perpendicular to the long axis of the mating pair was recorded. Profiles were background-subtracted and corrected for bleaching as follows: First, the fluorescence intensity was recorded over time in a small square on 12 non-mating cell and the mean was fitted to a double exponential:

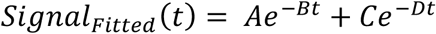

That fit was then used to divide the background-subtracted signal at each timepoint:

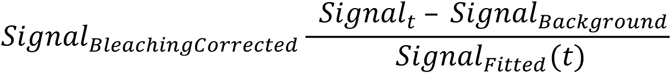

Corrected profiles were then either directly averaged and plotted, or further normalized to the mean of the WT maximum.

Patch to cytosol ratios were calculated from the ratio of the mean fluorescence intensity of 5 circular ROIs centered on patches to the mean fluorescence signal of 5 circular ROIs centered on cytosolic signal per cell. The same operation was repeated on 36 cells and plotted.

Total fluorescence intensities in mating pairs were obtained using single snapshots on regular slides, 7h post-starvation, by outlining the mating pairs and recoding the mean fluorescence intensity for each of them. Background fluorescence was assessed by a small square ROI in an area devoid of cells on each image, averaged over all images, and subtracted from fluorescence intensity measurements.

The number of ectopic foci was assessed using the 5-minutes time lapse overnight movies during the fusion process between the 2-dot stage and the fusion time by simply counting the number of time-frames showing an ectopic Myo52-tdTomato or any other marker. The numbers vary between experiments because the DeltaVision camera was upgraded in the middle of the project, allowing us to detect more delocalisation events. To assess colocalization, single and double-color ectopic foci were identified as above and the colocalization was calculated with the following formula:

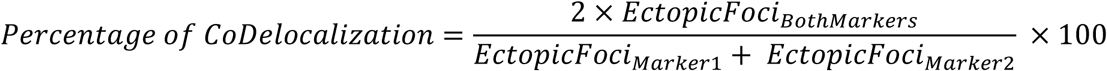

The proportion of ectopic foci containing a given marker was derived from the ratio of ectopic foci containing the given marker to the total number, which gave a ratio for each cell. That ratio was then averaged over all the recorded cells. For all experiments shown in Figure 6, only ectopic foci present at the cell sides were considered.

For3-3GFP retrograde flow was identified by visual inspection of spinning disk time-lapse imaging acquired at 1s interval. Only linear For3 dot movements present over at least 5 consecutive time frames were considered. Dots were manually tracked using the ImageJ multi-point tool and instantaneous speeds calculated and averaged per track. Kymographs were constructed using the ImageJ reslice tool along a 5-pixel-wide line along the For3 dot track.

All plots, fittings, corrections and normalisations were made using MATLAB home-made scripts. For boxplots, the central line indicates the median, the circle, if present, indicates the mean, and the bottom and top edges of the box indicate the 25th and 75th percentiles, respectively. The whiskers extend to the most extreme data points not considered outliers. For bar plots, errorbars represent the standard deviation. Statistical p-values were obtained using a two-sided t-test and are mentioned in the Figure legends, including sample size. For all the figures in this report, we consider, * < 5.10^−2^, ** < 5.10^−5^, *** < 5.10^−8^.

## Acknowledgements

We thank Dr Aleksandar Vjestica for plasmids, Dr Omaya Dudin for discussion and Dr Laura Merlini, Dr Olivia Muriel Lopez and Dr Veneta Gerganova for comments on the manuscript. This work was supported by an ERC Consolidator grant (CellFusion) and a Swiss National Science foundation grant (310030B_176396) to SGM.

## Supplemental material legends and Tables

**Movie S1. Formation of Myo52 ectopic focus and movement towards the fusion focus in *acp2*Δ.** Spinning-disk confocal microscopy time-lapse of Myo52-tdTomato and cytosolic GFP expressed under the *map3* promoter in WT and *acp2Δ* before fusion time. Time is in min:s. The movie is sped up 7x. Bars are 5µm.

**Movie S2. Coincidence of Fus1 and Myo52 at ectopic foci in *acp2*Δ.** Spinning-disk confocal microscopy time-lapse of Fus1-sfGFP and Myo52-tdTomato in WT and *acp2Δ* before fusion time. Time is in min:s. The movie is sped up 7x. Bars are 5µm.

**Movie S3. Localization of Fus1 ectopic foci at Fim1-labeled actin patches in *acp2*Δ.** Spinning-disk confocal microscopy time-lapse of Fus1-sfGFP and Fim1-mCherry in WT and *acp2Δ* before fusion time. Time is in min:s. The movie is sped up 7x. Bars are 5µm.

**Movie S4. Co-localization of tropomyosin Cdc8 and fimbrin in *acp2*Δ interphase cells.** Spinning-disk confocal microscopy time-lapse of GFP-Cdc8 and Fim1-mCherry in WT and *acp2Δ* in exponentially growing cells. Time is in min:s. The movie is sped up 7x. Bars are 5µm.

**Movie S5. Localization of For3 at Fim1-labeled actin patches in *acp2*Δ interphase cells.** Spinning-disk confocal microscopy time-lapse of For3-3GFP and Fim1-mCherry in WT and *acp2Δ* in exponentially growing cells. Time is in min:s. The movie is sped up 7x. Bars are 5µm.

**Table S1:**
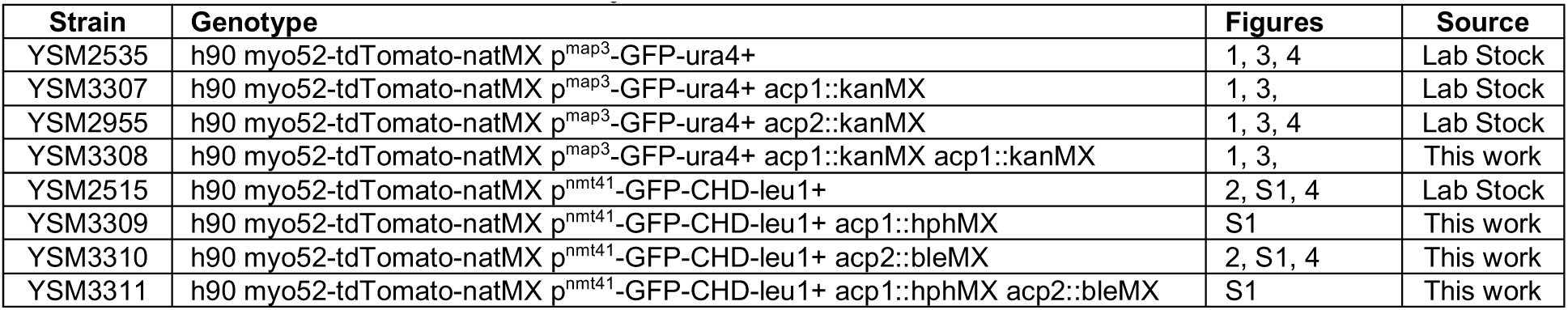
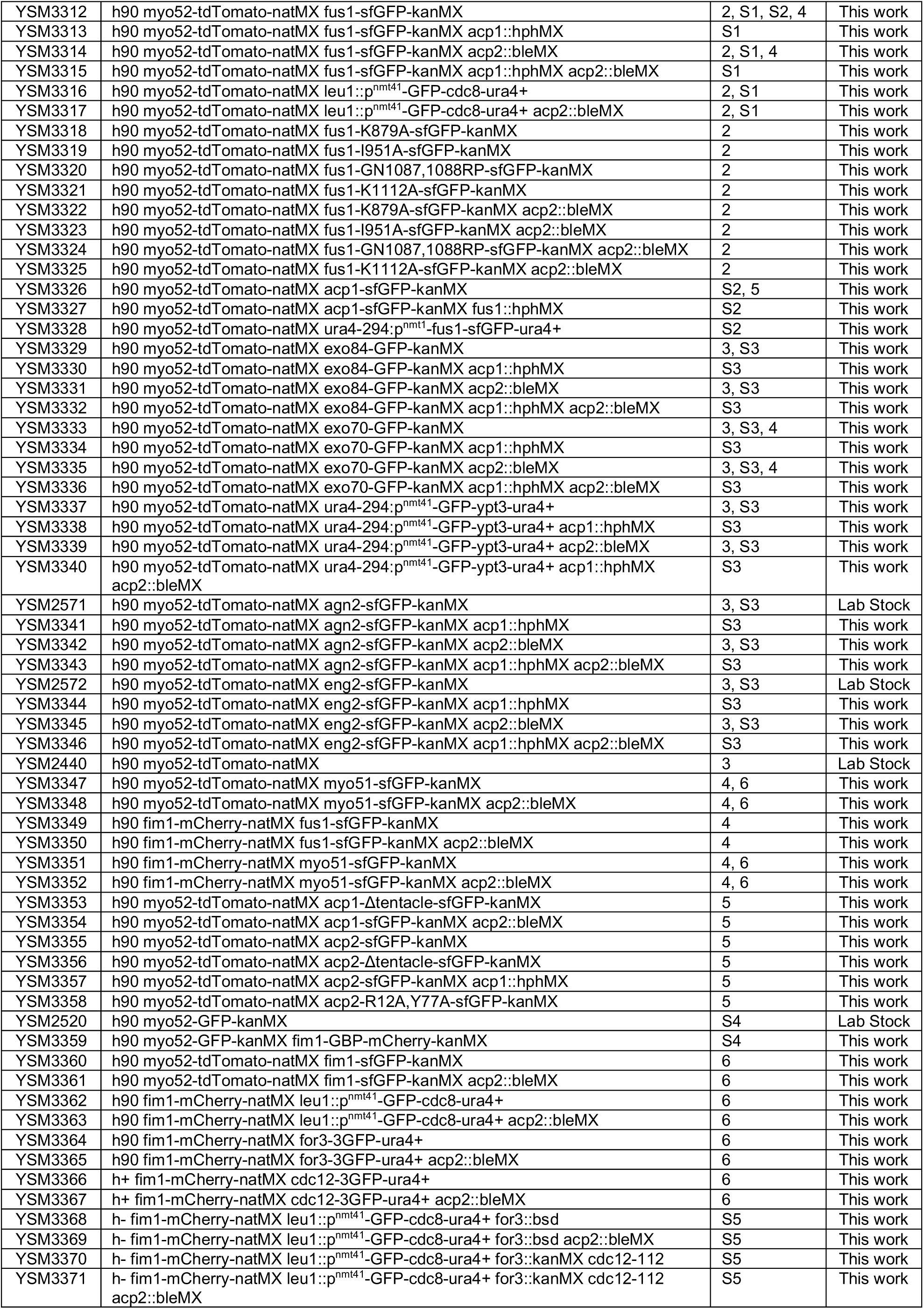
Strains used in this study.

